## Supplemental file for "NF45/NF90-mediated rDNA transcription provides a novel target for immunosuppressant development"

### Appendix

Appendix Figure S1

Appendix Figure S2

Appendix Figure S3

Appendix Figure S4

Appendix Figure S5

Appendix Figure S6

Appendix Figure S7

Appendix Figure S8

Appendix Table S1

Appendix Table S2

Appendix Table S3

**Appendix Fig S1**

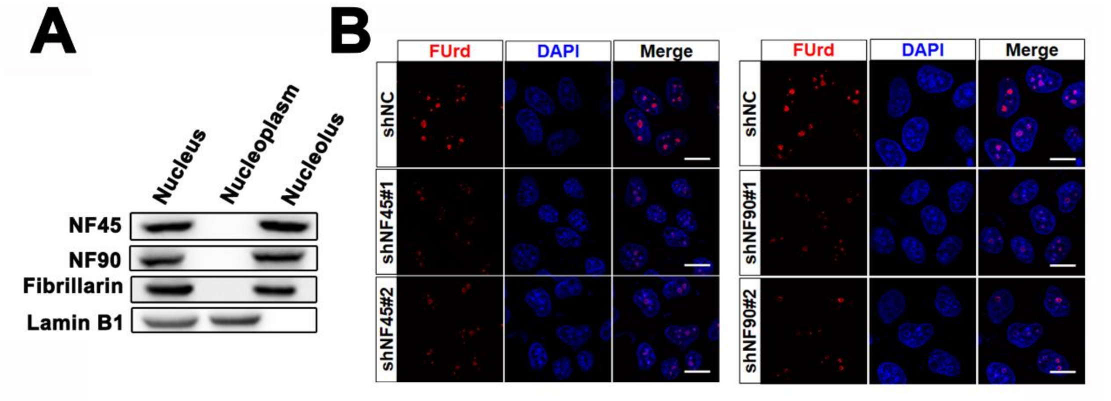


**Appendix Figure S1. NF45/NF90 is a nucleolar protein complex and positively regulates rDNA transcription** (related to Figure 1)**.**

A. HeLa cells were fractionated into nuclear, nucleoplasm and nucleolar fractions, and protein extracts were analyzed using western blot. Fibrillarin is a nucleolar marker; Lamin B1 is a nucleoplasm marker.

B. HeLa cells were silenced with negative control shRNA (shNC), NF45 shRNAs (shNF45#1 or sh4NF5#2) or NF90 shRNAs (shNF90#1 or shNF90#2), and then pulsed with FUrd for another 15 min in order to stain with an anti-BrdU (red). Scale bar, 10 $\mu$m.

**Appendix Fig S2**

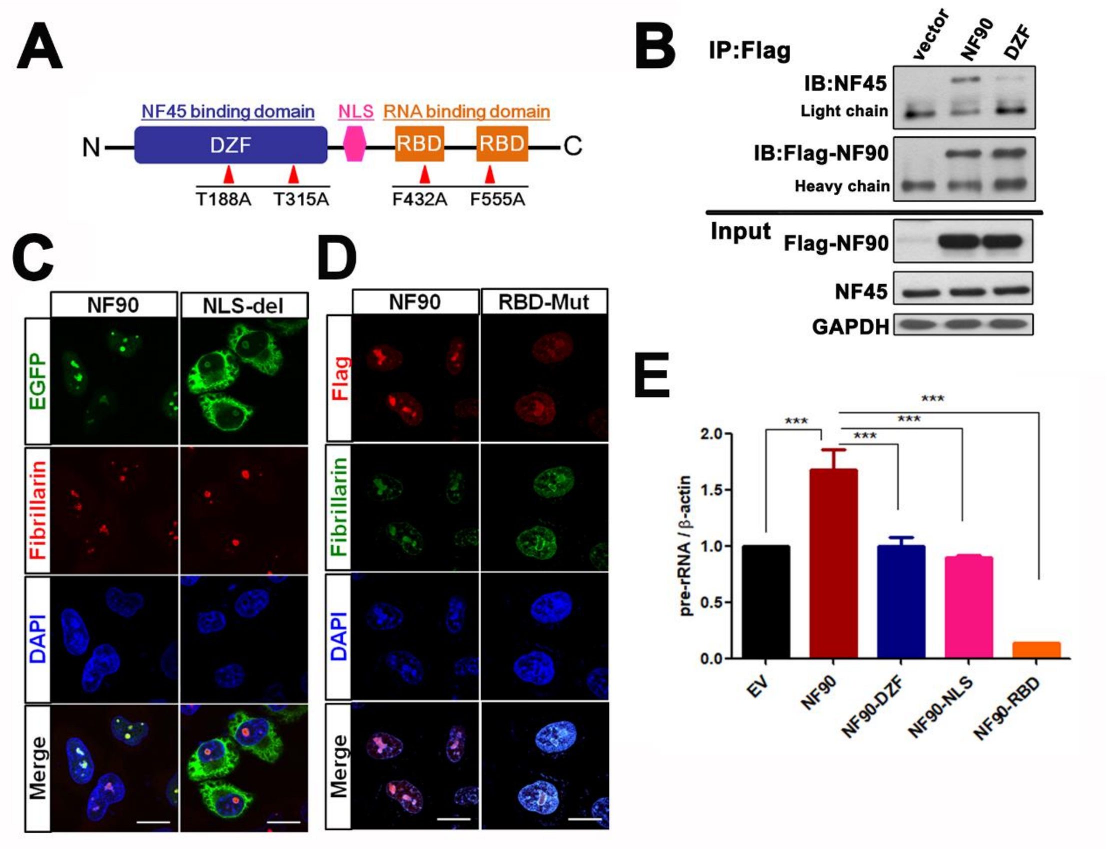


**Appendix Figure S2. The rDNA transcriptional regulation of NF45/NF90 requires them to act as a complex and locate in the nucleolus.** (related to Figure 1)**.**

A. Schematic diagram showing the domains of NF90 and its mutations (point mutations in DZF and RBD domains or NLS deleted mutation).

B. Co-IP analysis of NF45 with wild-type NF90 or DZF-mutated NF90.

C. Confocal images of HEK293 cells transfected with EGFP-NF90 or EGFP-NF90-NLS plasmid (fibrillarin is a nucleolar marker).

D. Confocal images of HEK293 cells transfected with FLAG-NF90 or FLAG-NF90-RBD plasmid. Scale bar, 10 $\mu$m.

E. HEK293 cells were transfected with an FLAG-NF90 vector or mutated vectors (DZF, NLS, or RBD) for 24 h, and then the total RNA was extracted from cells for qPCR analysis of pre-rRNA (n = 3) .

In all panels, Bar = mean ± SD. Statistical analysis by One-way ANOVA analysis followed by Tukey's test (E). (*** $p < 0.001$). Exact $p$ values are reported in Appendix Table S2.

**Appendix Fig S3**

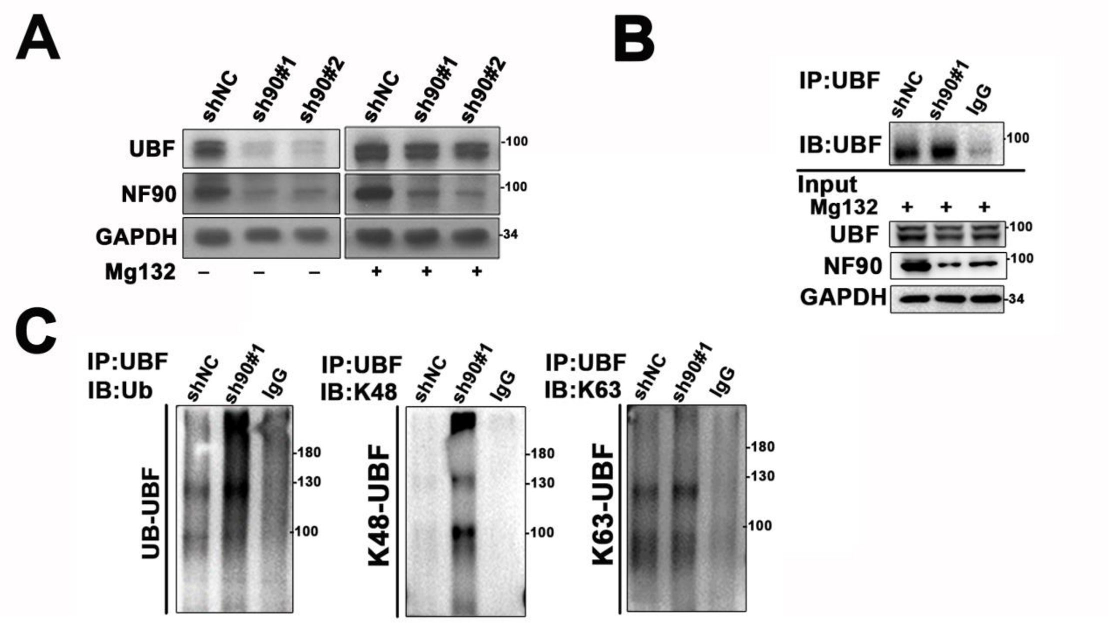


**Appendix Figure S3. NF90 knockdown increased UBF ubiquitination** (related to Figure 3)**.**

A. Western blot showing UBF and NF90 protein level in HEK293 cells infected with lentivirus containing NF90-specfic shRNAs (shNF90#1 and shNF90#2).

B. IP analysis of UBF in HEK293 infected with lentivirus containing shNF90#1.

C. IP analysis of UBF ubiquitination in HEK293 infected with lentivirus containing shNF90#1.

**Appendix Fig S4**

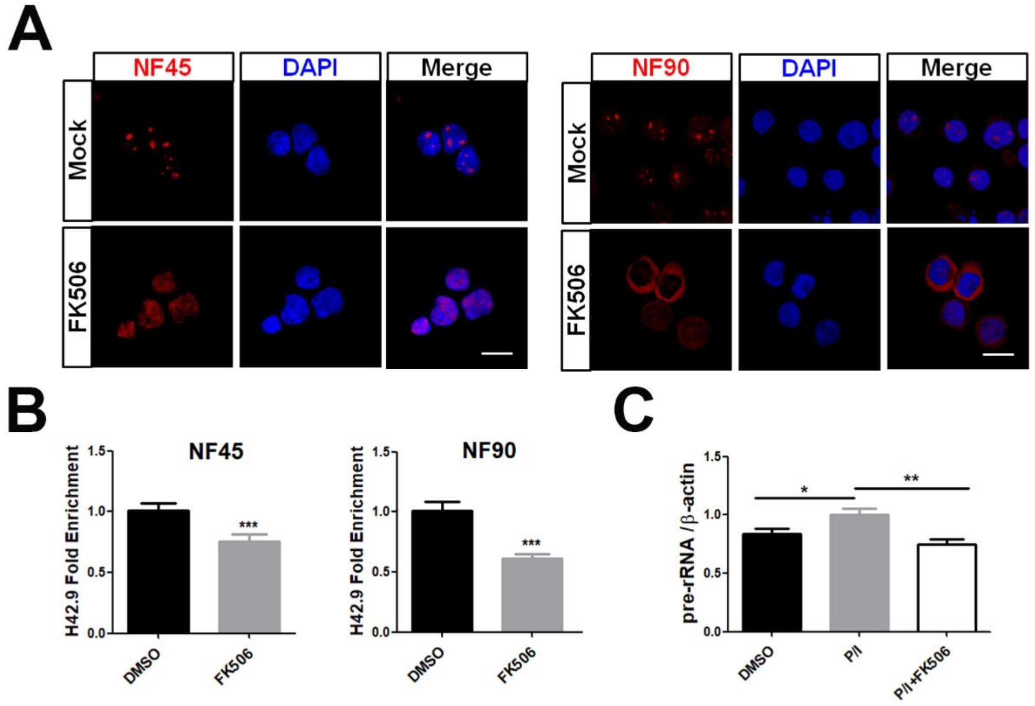


**Appendix Figure S4. FK506 affects the subcellular localization of NF45/NF90 and rDNA transcription in T cells** (related to Figure 4)**.**

A. Confocal images show the localization of NF45 (left panel) or NF90 (right panel) in P/I-stimulated $CD3^+$ T cells treated with or without FK506. Scale bar, 10 $\mu$m.

B. The binding of NF45 and NF90 to rDNA in P/I-stimulated $CD3^+$ T cells treated with or without FK506 (n = 3).

C. qPCR analysis of pre-rRNA in $CD3^+$ cells treated with P/I and a combination of P/I and FK506 (n = 3).

In all panels, Bar = mean ± SD. Statistical analysis by unpaired Student T-test (B). One-way ANOVA analysis followed by Tukey's test (C). (** $p < 0.01$, *** $p < 0.001$). Exact $p$ values are reported in Appendix Table S2.

**Appendix Fig S5**

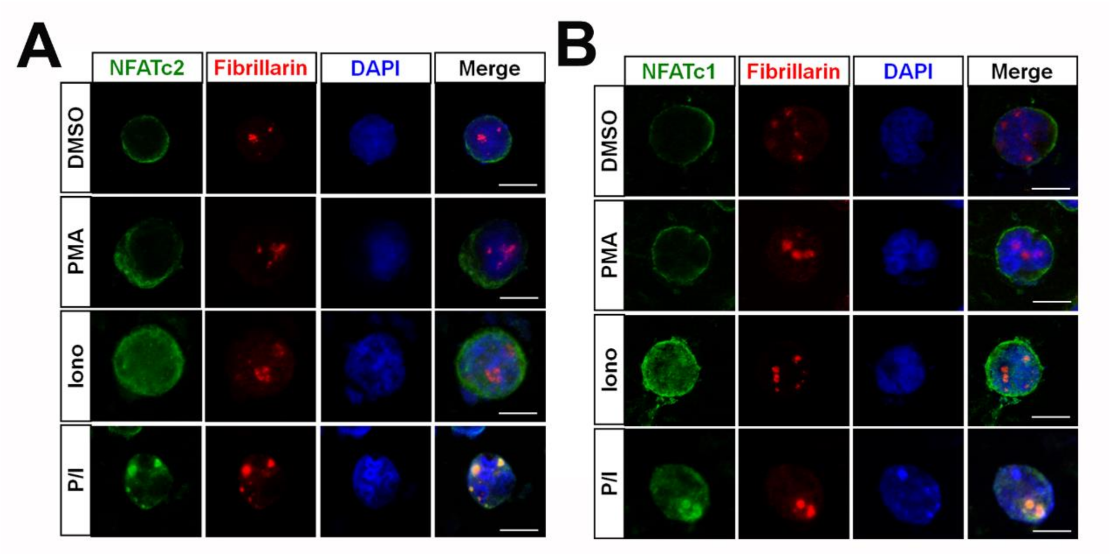


**Appendix Figure S5. NFATc2 and NFATc1 translocated to the nucleolus upon Iono or P/I stimulation** (related to Figure 4)**.**

A-B. Confocal images to show the subcellular localization of NFATc2 (A) or NFATc1 (B) in Jurkat cells treated with DMSO, PMA, Ionomycin or P/I. Scale bar, 10 $\mu$m.

**Appendix Fig S6**

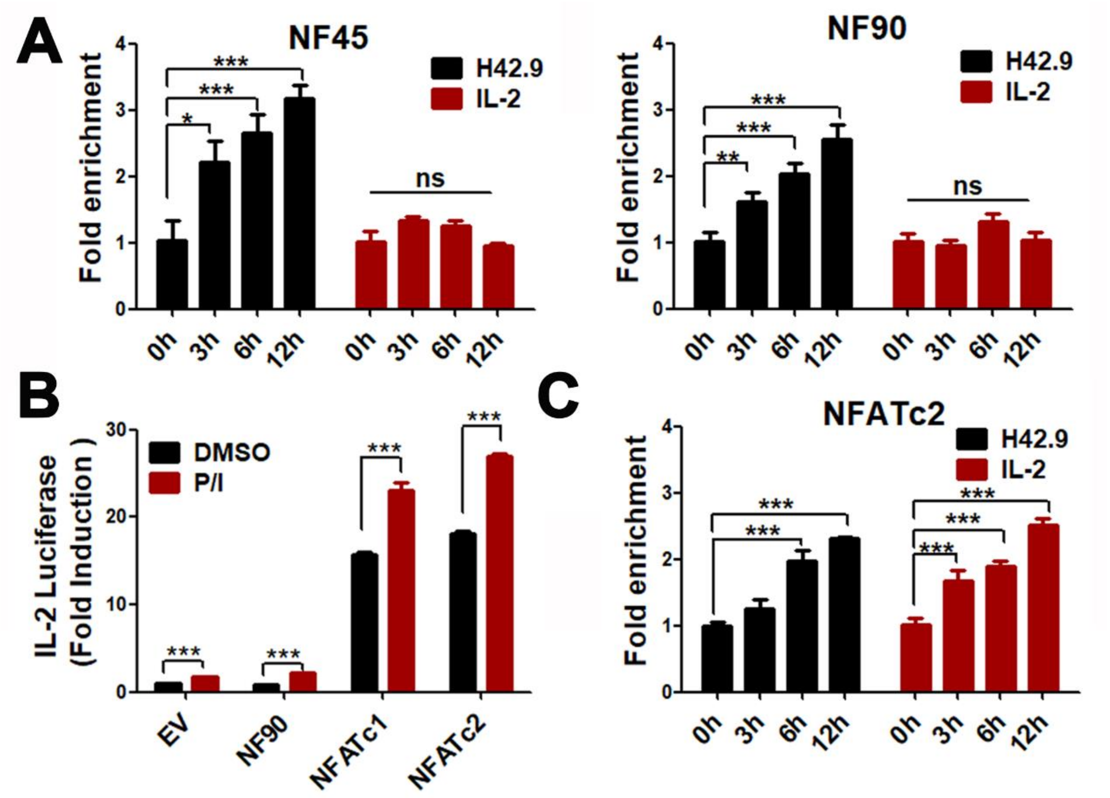


**Appendix Figure S6. The regulation functions of NF45/NF90 and NFATc1 or NFATc2 on rRNA transcription and IL-2 expression.**

A. The binding ability of NF45 or NF90 to the rDNA or IL-2 loci detected by a ChIP assay in $CD3^+$ cells stimulated with or without of P/I for the indicated times (n = 3).

B. Jurkat cells were transfected with empty vector (EV), FLAG-NF90, V5-NFATc2 or V5-NFATc1 in combination with IL-2-Luc reporter plasmids for 24 h, and then stimulated with or without of P/I for 12 h (n = 3) , The relative luciferase activity of IL-2 was detected.

C. The binding ability of NFATc2 to the rDNA or IL-2 loci detected by a ChIP assay in $CD3^+$ cells stimulated with or without of P/I for the indicated times (n = 3).

In all panels, Bar = mean ± SD. Statistical analysis by unpaired Student T-test (C). One-way ANOVA analysis followed by Tukey's test (A, B). (** $p < 0.01$, *** $p < 0.001$). Exact *p* values are reported in Appendix Table S2.

**Appendix Fig S7**

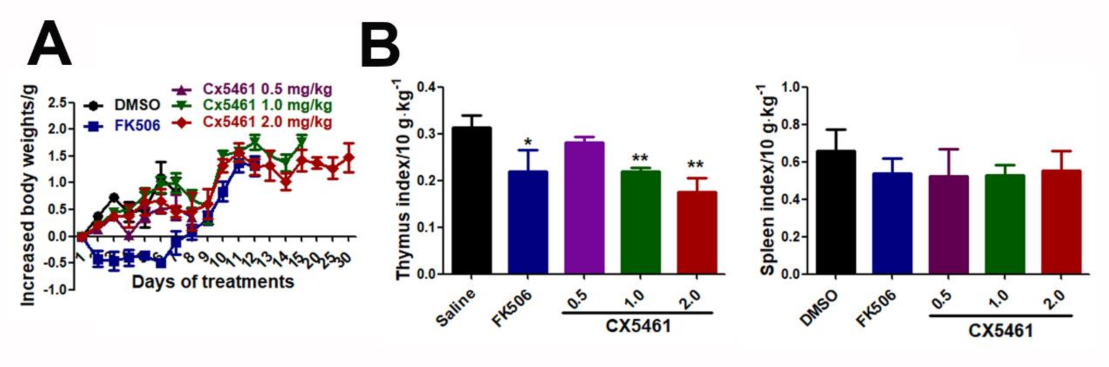


**Appendix Figure S7. The effects of CX5461 on the body weight, spleen weight and thymus weight of skin-graft mice** (related to Figure 6)**.**

A. Plots of the body weight of skin-graft mice after injecting with DMSO, FK506 or different doses of CX5461.

B. The thymuses and spleens of each group were weighed at the end point of the study (n = 5). In all panels, Bar = mean ± SD. Statistical analysis by One-way ANOVA analysis followed by Tukey’s test (B). (* $p < 0.05$, *** $p < 0.001$). Exact *p* values are reported in Appendix Table S2.

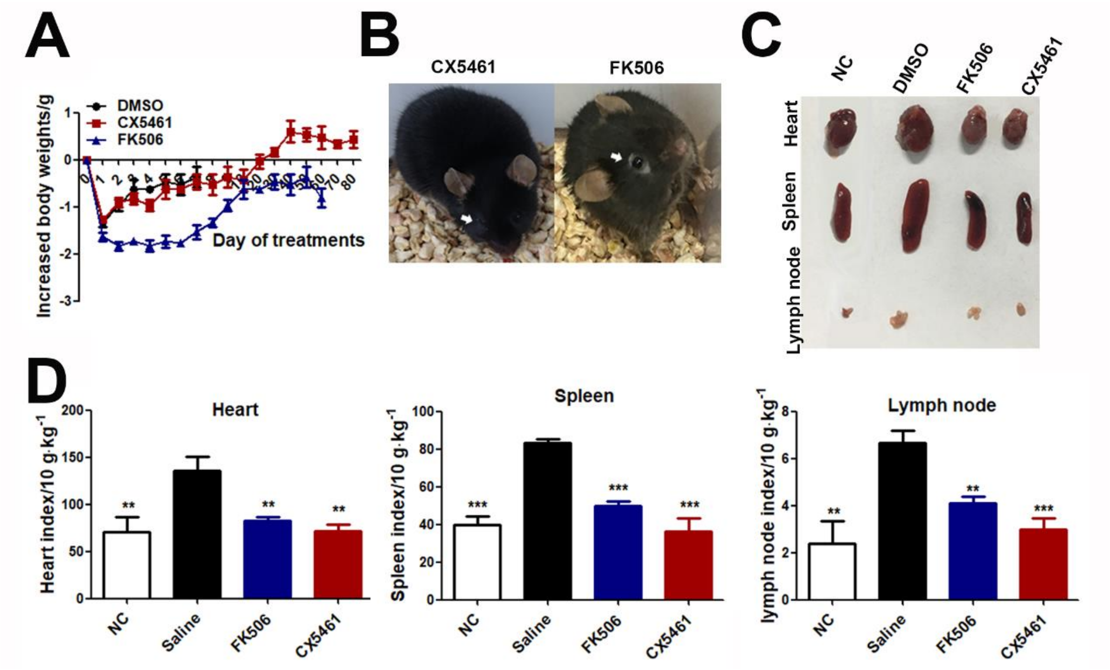


**Appendix Figure S8. The effects of CX5461 on the body weight, hair loss, and spleen and thymus weights of heart-transplant mice** (related to Figure 6)**.**

A. Plots of body weights of heart-transplant mice after injecting with DMSO, FK506 or CX5461.

B. Images highlighting the hair loss of heart-transplant mice after injecting with FK506 or CX5461 at Day 7.

C-D. Images (C) and weights (D) of the viscera of heart-grafts, spleens and lymph nodes of each group at the end point of the study (n = 5).

In all panels, Bar = mean ± SD. Statistical analysis by One-way ANOVA analysis followed by Tukey's test (D). (** $p < 0.01$, *** $p < 0.001$). Exact *p* values are reported in Appendix Table S2.

**Appendix Table S1. The demographic and baseline clinical characteristics of the study in kidney transplant patients**

| Variables | Total (n = 23) | ABMR (n = 8) | TCMR (n = 8) | stable (n = 7) | *P* |
|---|---|---|---|---|---|
| Donors | | | | | |
| Types, n (%) | | | | | **0.005** |
| DD | 13 (57) | 2 (25) | 8 (100) | 3 (43) | |
| LD | 10 (43) | 6 (75) | 0 (0) | 4 (57) | |
| Recipients | | | | | |
| Gender, n (%) | | | | | **0.017** |
| Female | 9 (39) | 1 (12) | 2 (25) | 6 (86) | |
| Male | 14 (61) | 7 (88) | 6 (75) | 1 (14) | |
| Age (year), Median (IQR) | 33.81 (31.40, 39.65) | 37.84 (34.13, 48.32) | 32.97 (30.31, 41.70) | 31.72 (30.76, 33.50) | 0.102 |
| Weight (kg), Median (IQR) | 52.0 (43.8, 63.8) | 63.3 (52.0, 64.0) | 54.3 (44.9, 63.3) | 42.5 (41.4, 50.0) | 0.138 |
| Previous transplantation, n (%) | | | | | 0.494 |
| 0 | 20 (87) | 8 (100) | 6 (75) | 6 (86) | |
| 1 | 3 (13) | 0 (0) | 2 (25) | 1 (14) | |
| Days from transplant to sample, Median (IQR) | 366 (140, 582) | 1474 (633, 2353) | 128 (103, 149) | 376 (369, 378) | **0.001** |
| CNIs [*], n (%) | | | | | 1.000 |
| CsA | 1 (4) | 0 (0) | 1 (12) | 0 (0) | |
| FK506 | 22 (96) | 8 (100) | 7 (88) | 7 (100) | |
| FK506 $C_0$ (ng/ml), Median (IQR) | 6.6 (5.3, 7.8) | 7.7 (7.2, 8.5) | 5.7 (5.1, 6.6) | 6.3 (5.6, 8.3) | 0.161 |
| CsA $C_0$ (ug/L) [#] | — | — | 127.5 | — | — |
| White blood cell count (10^9/L), Median (IQR) | 6.99 (5.98, 9.21) | 7.00 (6.22, 8.97) | 7.44 (6.39, 9.86) | 6.41 (5.98, 8.25) | 0.694 |
| Lymphocyte count (10^9/L), Mean ± SD | 1.41 ± 0.68 | 1.13 ± 0.37 | 1.53 ± 0.82 | 1.61 ± 0.75 | 0.339 |
| Neutrophil (10^9/L), Median (IQR) | 4.65 (3.74, 6.85) | 5.05 (4.30, 7.62) | 6.06 (4.29, 6.87) | 3.80 (3.54, 5.65) | 0.408 |
| Monocyte count (10^9/L), Mean ± SD | 0.61 ± 0.21 | 0.62 ± 0.18 | 0.68 ± 0.29 | 0.52 ± 0.14 | 0.363 |

CNIs, Calcineurin inhibitors; FK506, Tacrolimus; CsA, Cyclosporin A; LD, Living donor; DD, Death donor. * Standard maintenance immunosuppression, including CNI, prednisolone and mycophenolic acid was applied to all patients. # Only one patient received cyclosporine A, and no standard deviation of CsA $C_0$ was provided.

### **Appendix Table S3. List of** qPCR primers

| **Gene** | **Forward primer sequence 5’→3’** | **Reverse primer sequence 5’→3’** |
|---|---|---|
| Human *Actb* | **GAACGGTGGTGTGTCGTTC** | **GCGTCTCGTCTCGTCTCACT** |
| Human pre-rRNA | **GAACGGTGGTGTGTCGTTC** | **GCGTCTCGTCTCGTCACT** |
| Mouse *Actb* | **GGCTGTATTCCCCTCCATCG** | **CCAGTTGGTAACAATGCCATGT** |
| Mouse pre-rRNA | **CTCCTGTCTGTGGTGTCCAA** | **GCTGGCAGAACGAGAAGAAC** |
| Human H0 | **GGTATATCTTTCGCTCCGAG** | **GACGACAGGTCGCCAGAGGA** |
| Human H13 | **ACCTGGCGCTAAACCATTCGT** | **GGACAAACCCTTGTGTCGAGG** |
| Human H23 | **CCTTCCACGAGAGTGAGAAGC** | **TCGACCTCCCGAAATCGTACA** |
| Human H42.9 | **CCCGGGGGAGGTATATCTTT** | **CCAACCTCTCCGACGACA** |
| Human IL-2 promoter | **TAAGTGTGGGCTAACCCGA** | **AAGGAGCACAAGTGTCAATGTGA** |

**Appendix Table S3. List of exact *p*-values**

| Figure | Panel | Sub-pancel | Compared groups | *P*-value |
|---|---|---|---|---|
| Figure 1 | B | shNF45 | shNF45#1 vs NC | 0.0202 |
| | | | shNF45#2 vs NC | 0.0125 |
| | | shNF90 | shNF90#1 vs NC | <0.001 |
| | | | shNF90#2 vs NC | <0.001 |
| | C | NF45 | NF45 vs EV | <0.001 |
| | | NF90 | NF90 vs EV | <0.001 |
| | E | shNF45 | shNF45#1 vs NC | <0.001 |
| | | | shNF45#2 vs NC | <0.001 |
| | | shNF90 | shNF90#1 vs NC | <0.001 |
| | | | shNF90#2 vs NC | <0.001 |
| | F | NF45 | NF45 vs EV | <0.001 |
| | | NF90 | NF90 vs EV | <0.001 |
| Figure 2 | E | NF90- | Mutant vs WT | <0.001 |
| | | NF90+ | Mutant vs WT | <0.001 |
| | | WT | NF90+ vs NF90- | <0.001 |
| Figure 3 | E | NF45 | 3 h vs 0 h | 0.0060 |
| | | | 6 h vs 0 h | <0.001 |
| | | NF90 | 3 h vs 0 h | 0.0053 |
| | | | 6 h vs 0 h | <0.001 |
| | F | NF45 | NF45 vs EV | <0.001 |
| | | NF90 | NF90 vs EV | <0.001 |
| Figure 4 | A | NF45 | P/I vs DMSO | 0.0283 |
| | | NF90 | P/I vs DMSO | <0.001 |
| | B | | P/I vs DMSO | 0.0177 |
| | C | shNC | P/I vs DMSO | <0.001 |
| | D | shNC | P/I vs DMSO | <0.001 |
| | G | DMSO | NFATc2 vs IgG | <0.001 |
| | | P/I | NFATc2 vs IgG | <0.001 |
| | H | WT | EV vs NF90 | <0.001 |
| | | PI | NF90 vs EV | <0.001 |
| | | | NFATc2 vs EV | <0.001 |
| | | | NF90+NFATc2 vs NF90 | <0.001 |
| | | | NF90+NFATc2 vs NFAT | <0.001 |
| | I | shNC | P/I vs DMSO | <0.001 |
| Figure 5 | B | | shNF45#1 vs NC | <0.001 |
| | | | shNF45#2 vs NC | <0.001 |
| | | | shNF90#1 vs NC | <0.001 |
| | | | shNF90#2 vs NC | <0.001 |
| | C | | shNF45#1 vs NC | 0.0278 |
| | | | shNF45#2 vs NC | 0.0147 |

| | | | | |
|---|---|---|---|---|
| | | | shNF90#1 vs NC<br>shNF90#2 vs NC | 0.0027<br>0.0113 |
| | G | | DMSO vs P/I<br>P/I+FK506 vs P/I<br>P/I+CX5461 vs P/I | <0.001<br><0.001<br><0.001 |
| | H | | DMSO vs P/I<br>P/I+FK506 vs P/I<br>P/I+CX5461 vs P/I | <0.001<br>0.0366<br><0.001 |
| Figure 6 | A | Skin allograft<br>Heart allograft | Rejection vs NC<br>Rejection vs NC | <0.001<br><0.001 |
| | B | | TCMR vs Stable | 0.0042 |
| | G | | FK506 vs DMSO<br>CX54610 1.0 vs DMSO<br>CX54610 2.0 vs DMSO | 0.0267<br>0.0113<br>0.0061 |
| | H | | FK506 vs DMSO<br>CX54610 0.5 vs DMSO<br>CX54610 1.0 vs DMSO<br>CX54610 2.0 vs DMSO | 0.0069<br>0.0197<br>0.0035<br>0.0036 |
| | I | | FK506 vs DMSO<br>CX5461 vs DMSO | 0.0099<br><0.001 |
| | L | | NC vs DMSO<br>FK506 vs DMSO<br>CX5461 vs DMSO | <0.001<br><0.001<br><0.001 |
| | M | | NC vs DMSO<br>FK506 vs DMSO<br>CX5461 vs DMSO | <0.001<br>0.0070<br><0.001 |
| | N | | NC vs DMSO<br>FK506 vs DMSO<br>CX5461 vs DMSO | <0.001<br><0.001<br><0.001 |
| Appendix Figure S2 | E | | NF90 vs EV<br>NF90-DZF vs EV<br>NF90-NLS vs EV<br>NF90-RBD vs EV | <0.001<br><0.001<br><0.001<br><0.001 |
| Appendix Figure S4 | B | NF45 | FK506 vs DMSO | <0.001 |
| | | NF90 | FK506 vs DMSO | <0.001 |
| | C | | DMSO vs P/I<br>P/I+FK506 vs P/I | 0.0177<br>0.0034 |
| Appendix Figure S6 | A | NF45- H42.9 | 3 h vs 0 h | 0.0140 |
| | | | 6 h vs 0 h | <0.001 |
| | | | 12 h vs 0 h | <0.001 |
| | | NF90- H42.9 | 3 h vs 0 h | 0.0056 |
| | | | 6 h vs 0 h | <0.001 |
| | | | 12 h vs 0 h | <0.001 |

| | | | | |
|---|---|---|---|---|
| | B | NFATc2- H42.9 | 6 h vs 0 h | <0.001 |
| | | | 12 h vs 0 h | <0.001 |
| | | NFATc2- IL-2 | 3 h vs 0 h | <0.001 |
| | | | 6 h vs 0 h | <0.001 |
| | | | 12h vs 0h | <0.001 |
| | C | EV | PI vs DMSO | <0.001 |
| | | NF90 | PI vs DMSO | <0.001 |
| | | NFATc1 | PI vs DMSO | <0.001 |
| | | NFATc2 | PI vs DMSO | <0.001 |
| Appendix Figure S7 | B | Thymus index | FK506 vs DMSO<br>CX5461 1.0 vs DMSO<br>CX5461 2.0 vs DMSO | 0.0378<br>0.0038<br>0.0039 |
| Appendix Figure S8 | D | Heart | NC vs DMSO<br>FK506 vs DMSO<br>CX5461 vs DMSO | 0.0070<br>0.0049<br>0.0031 |
| | | Spleen | NC vs DMSO<br>FK506 vs DMSO<br>CX5461 vs DMSO | <0.001<br><0.001<br><0.001 |
| | | Lymph node | NC vs DMSO<br>FK506 vs DMSO<br>CX5461 vs DMSO | 0.0026<br>0.0021<br><0.001 |
